# Emergence and patterning dynamics of mouse definitive endoderm

**DOI:** 10.1101/728642

**Authors:** Maayan Pour, Abhishek Sampath Kumar, Maria Walther, Lars Wittler, Alexander Meissner, Iftach Nachman

## Abstract

The segregation of definitive endoderm (DE) from mesendoderm progenitors leads to the formation of two distinct germ layers. Dissecting DE onset has been challenging as it occurs within a narrow spatio-temporal window in the embryo. Here we employ a dual Bra-GFP, Sox17-RFP reporter cell line to study DE onset dynamics. We find Sox17 starts in a few isolated cells *in vivo*. Using 2D and 3D *in vitro* models, we show that DE cells emerge from mesendoderm progenitors at a temporally regular, but spatially stochastic pattern, which is subsequently arranged by self-sorting of Sox17+ cells. Self-sorting coincides with up-regulation of E-cadherin but is not necessary for DE differentiation or proliferation. A subpopulation of Bra-high cells commits to a Sox17+ fate independent of external Wnt signal. Our *in vivo* and *in vitro* results highlight basic rules governing DE onset and patterning through the commonalities and differences between these systems.

## Introduction

Early in mouse embryonic development, epiblast cells undergo gastrulation, the process that derives the three primary germ layers. Bi-potent mesendoderm cells egress through the extending primitive streak and segregate to two distinct lineages of the body, mesoderm and definitive endoderm (DE) [1] The decision of mesendoderm progenitor cells to become DE is thus one of the earliest developmental decisions during embryogenesis. Ingressing epiblast cells switch on their DE identity while moving ventrally and laterally, eventually intercalating with the visceral endoderm layer. In this process they segregate from future mesodermal cells, eventually resulting in distinct germ layers with the correct ratio of cell numbers. Several signaling pathways, such as Wnt and Nodal/Activin, are known to play dominant roles in driving this decision [1–4]. WNT signaling plays an essential function in mesendoderm specification by controlling the expression of Nodal and its co-receptor Cripto during gastrulation [1, 5]. Mesendoderm progenitors initially start from a Wnt-high region (the proximal-posterior PS), and future DE cells migrate toward the anterior side, a Nodal-high region [6, 7]. It is not clear whether local signaling gradients or stochastic epigenetic or expression state differences drive the decision of specific cells to become DE, nor at what point cells commit to their DE fate.

One of the challenges to study DE decision in the mouse embryo is the lack of unique markers, as most of the endoderm markers are also expressed in the primitive endoderm (PrE) or visceral endoderm (VE). For example, Sox17, one of the earliest DE regulators, is expressed in VE (PrE), DE, and later endodermal tissues in the developing embryo [8, 9]. Recently it has been shown that both epiblast-derived DE cells and visceral endoderm (VE) cells converge not only spatially (through egression and intercalation), but also transcriptionally, both contributing to endodermal structures, such as the gut endoderm [10–12]. Therefore, in order to decipher the onset and dynamics of the epiblast-derived DE, there is a need to selectively track these cells and distinguish them from extra-embryonic derived endoderm. Even then, it is hard to tease apart the contribution of other crucial factors such as cellular localization, signaling gradient and expression profile, as these are all coupled in the developing embryo. In-vitro 2D and 3D systems have been used in recent years to model early stages of embryonic development, including mesendoderm progression [13–20]. These provide accessible systems, amenable to manipulation and decoupling of different external conditions. The comparative view of commonalities between such systems and the embryo provide a promising opportunity to dissect basic rules of cell fate decisions and early patterning [21].

Here we set out to study the onset dynamics and progression of the epiblast-derived Sox17+ cell population (DE), using a newly engineered dual Bra-GFP, Sox17-RFP reporter cell line. Double positive embryos derived through tetraploid embryo complementation experiments reveal that the DE emerges in few isolated cells within the mesendoderm spread population. To better understand the factors driving the DE fate decision, we dissected its dynamics in complementary 2D and 3D *in vitro* contexts. In both systems, DE cells arise from within the mesendoderm population in a temporally-synchronized manner, but in a spatially random “salt-and-pepper” pattern. This is followed by a self-sorting phase of the Sox17+ cells, leading to aggregation (2D) or lumenogenesis (3D) as the in-vitro counterpart to intercalation in the embryo. This self-sorting precedes high E-cadherin expression, and is not essential for DE differentiation or proliferation. Analyzing the temporal dependencies of DE differentiation on Wnt and Nodal signaling, we find that a small subpopulation of high-expressing Bra cells are already committed to their Sox17+ fate independent of external Wnt signal. The robust properties we observe in the different *in vitro* contexts shed new light on principles underlying DE dynamics in-vivo.

## Results

### Sox17 onsets in the primitive streak in few isolated cells in vivo

To study the onset of mesendoderm derived DE and the relative patterning of DE and mesendoderm progenitors, we have created a dual-reporter Sox17-RFP (mStrawberry), Brachyury-GFP (Bra-GFP) mESC line (Fig. 1a, b, Fig. S1). We then generated transgenic mouse embryos from the reporter cells by tetraploid complementation such that all embryonic tissues (e.g. primitive streak and mesendoderm progenitors) are derived from the diploid transgenic mESC line, while extra-embryonic tissues (including visceral endoderm) are from the wild type tetraploid donor. We imaged these embryos between gastrulation initiation (E6.5) and early organogenesis (E8.5) to monitor the onset and expansion pattern of Sox17-RFP within the embryonic tissues. In gastrulating embryos, the first Sox17-RFP+ cells arise within the Bra-GFP+ cells at the anterior part of the PS (Fig. 1c). Unlike the Bra-GFP+ cell population which is contiguous, Sox17 initially arises in a few isolated cells (Fig. 1d). Consistent with previous observations, the Sox17+ population expands during migration laterally toward the anterior side, and is spread over the whole distal side of the epiblast by E7.5 [10]. These cells later intercalate with the cells of the visceral endoderm and undergo epithelialization, and by E8.5 all embryo-derived Sox17+ cells are localized along the gut tube (Fig. 1c). These observations raise questions about how the initial DE population is specified, and what mechanisms set its resulting pattern.

**Figure 1.**
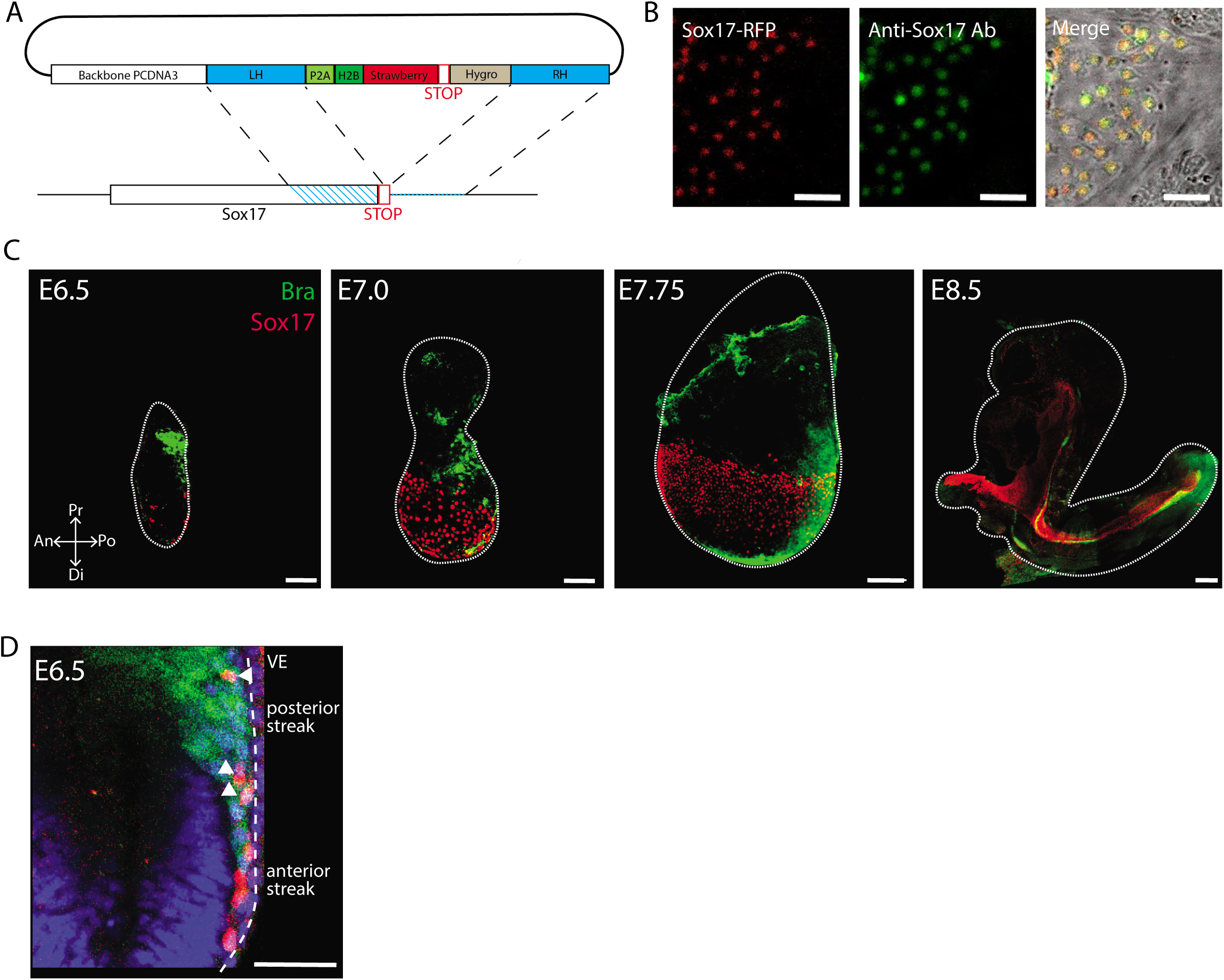
Sox17 emerges in few isolated cells in the mouse tetraploid embryos. (A) CRISPR knock-in of the H2B-RFP cassette in the endogenous Sox17 gene design. (B) Overlap between Sox17-RFP live marker (left) and Sox17 antibody immunostaining (middle) to validate the live marker cell line in a 2D colony. Scale bar 50um. (C) Images of Maximum intensity projections of E6.5-E8.5 tetraploid embryos with E14 Bra-GFP and Sox17-RFP double marker cell line, shows the expression and resolution of Bra+/Sox17+ cells in the gastrulating embryos. Dashed line marks the embryo borders. Scale bar 100um. (D) Single Z-slice image within E6.5 Embryo shows the emergence of Sox17+ cells within the Bra+ cells in the primitive streak. Scale bar 50 um.

### Sox17 emerges from Bra expressing cells in a fixed temporal pattern

To better understand the rules governing onset and expansion of Sox17 within mesendoderm cells, we differentiated our reporter ES cells within 2D colonies or 3D embryoid bodies (EBs) in defined medium (N2B27). At 48 hours, CHIR (a Wnt signaling activator) was added, as these conditions have been shown before to induce both mesoderm and endoderm [14, 15]. We first verified the identity and origin of Sox17-RFP+ cells in the 2D and 3D cultures and did not observe RFP+ cells under pluripotency conditions, except for peripheral cells in 2D colonies, but not in 3D embryoid bodies (Fig. S2). We next tracked the origin of DE cells in the 2D and 3D cultures. Bra expression precedes Sox17 at the RNA level during EB differentiation (Fig. S3). To have a better temporal resolution of this dynamic, the GFP and RFP signals were quantified over time in each EB/colony. We designated the time of CHIR addition as 0 hrs (Fig. 2a) and found that in embryoid bodies, Sox17 emerged at a highly uniform temporal window from Brachyury positive cells (Fig. 2b). The first Sox17-RFP cells are observed 24±1.5 hours (n=13) after addition of CHIR, 13±4 hours after the onset of Brachyury, and 2±1.5 hours before the peak of Brachyury expansion. This pattern is highly reproducible in individual EBs, with high correlation between the onset of the two genes (Fig. 2d,e, S4), suggesting they are part of a sequential differentiation process. Interestingly, in 2D colonies the dynamics is delayed with Sox17 emerging at a later time (37±4 hrs), approximately 20±3 hours after Bra onset, but with a close proximity to Bra peak (39±2 hrs). While the expression dynamics of both Bra and Sox17 in 2D is longer then in 3D, the relative temporal pattern is similar (Fig. 2c). The robustness of the relative pattern to the differences between the 2D and 3D setups suggests this sequential process is not dependent on cell interactions with the environment or other cells.

**Figure 2.**
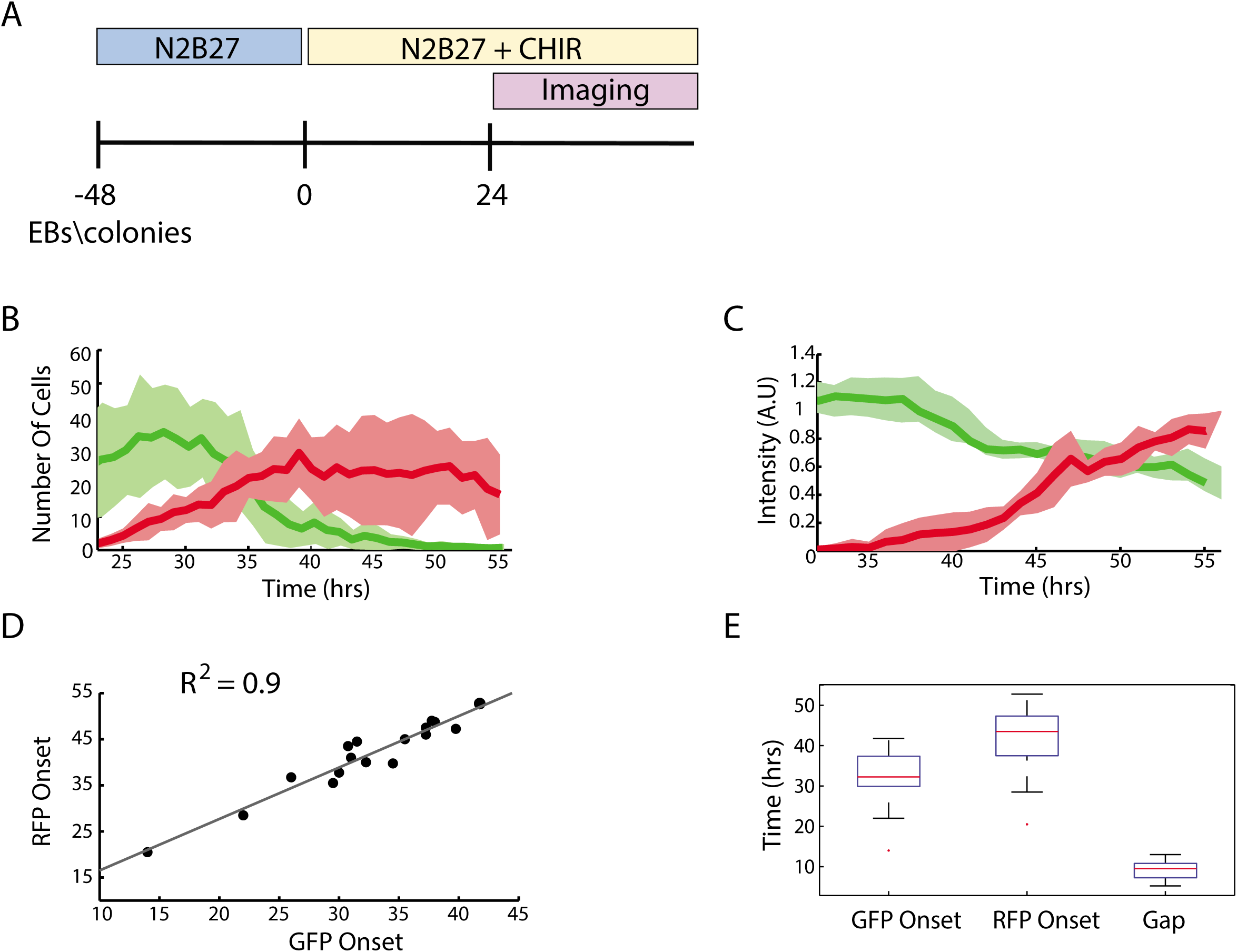
A fixed temporal pattern between Brachyury and Sox17 onsets. (A) Experiment design. ESCs were grown in N2B27 medium as 2D colonies or as EBs. After 48 hours, CHIR was added to the medium (marked as 0 hrs). Imaging started at 24 hours. (B) Number of Bra-GFP (green) cells or number of Sox17-RFP (red) cells per EB over time. Mean and SD of 10 EBs are shown. (C) Total intensity of Bra-GFP (green) or Sox17-RFP (red) per 2D colony over time. Mean and SD of 10 colonies are shown. (D) Bra-GFP Onset time versus Sox17-RFP Onset time in EBs. Each dot represents one EB (n=18). (E) Boxplot of Bra-GFP onset time, Sox17-RFP onset time and the gap between them.

### Sox17 emerges in a salt-and-pepper pattern, followed by self-sorting

We next wanted to characterize the spatial pattern of Sox17 emergence within the Brachyury-positive population. In embryoid bodies, Sox17-RFP+ cells initially emerge within the Bra-GFP+ population, close to the peak of Bra expression. The spatial pattern of appearance is salt-and-pepper like: RFP+ cells are randomly distributed within the GFP+ population, with no clear spatial organization, similar to their early pattern in the embryo (Fig. 1c). However, the cells begin to self-sort 13±1.5 hours after the appearance of Sox17, moving within the GFP+ population, eventually finding each other to form either internal lumens or an envelope at the outer shell of the EB (Fig. 3a, Movie S1). A similar spatial dynamic is observed in 2D colonies (Fig. 3b, Movie S2), where after sorting out of the GFP+ area, the Sox17-RFP+ cells form distinct continuous patches. To quantify the amount of spatial bias at the onset of Sox17, we analyzed the distribution of distances between all pairs of Sox17-RFP+ cells (Fig. 3c,d). At the onset time (when we spot 7-10 RFP+ cells within the EB), this distribution is indistinguishable from that of all cells in the EB volume (p=0.8, two-sample Kolmogorov-Smirnov test), consistent with random locations and no spatial bias (Fig. 3c). In contrast, after 24-30 hours, when RFP+ cells have self-sorted into consistent lumens/aggregates, the distance distribution of RFP+ cells significantly differs from that of all cells (p=4E-27) (Fig. 3d). To quantify the aggregation dynamics of the RFP+ cells in EBs, we performed a coarse segmentation, to count the number of continuous cell clusters (Fig. 3e, Movie S3). For example, each formed lumen is counted as a single cluster. Initially, the number of clusters increases with the number of cells. Once self-sorting begins, the number of clusters begins to decrease, as Sox17-RFP+ cells or clusters find each other and merge into larger clusters.

**Figure 3.**
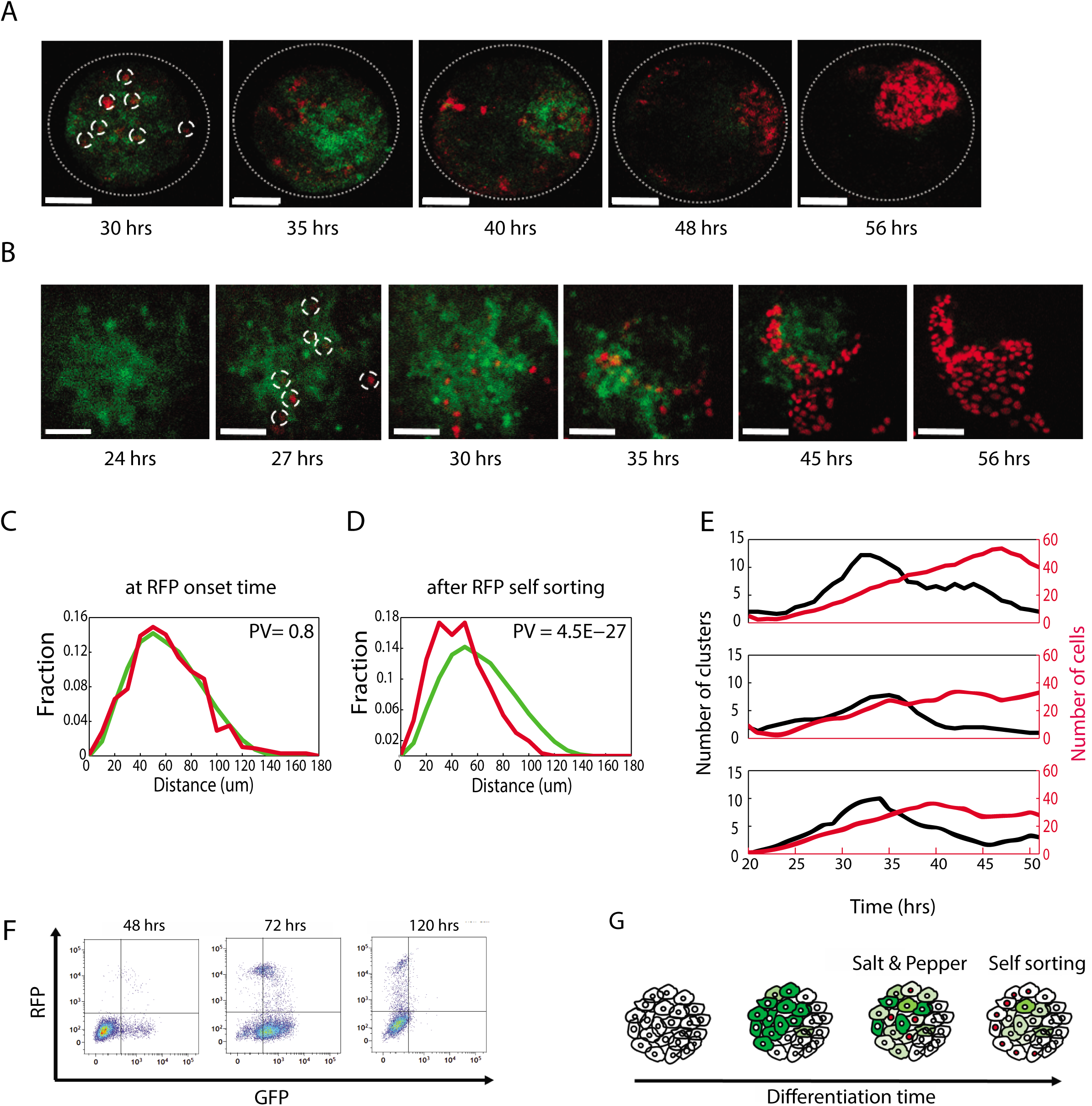
Sox17 onsets in a salt-and-pepper pattern followed by self-sorting. (A,B) Sox17-RFP cells emerge within the Bra-GFP population and expand to form an RFP only locus in EBs (A) or in 2D colonies (B). Scale bar 50um. (C,D) The distribution of distances between RFP+ cells (red) compared to distances between all cells (green). The two distributions are similar at the time or sox17 onset (P = 0.8151, Kolmogorov-Smirnov test) (C). At 56 hours of differentiation, the two distributions differ significantly (P= 4.4617E-27), after RFP cells have clustered together (D). (E) Number of sox17 cells (red line, right Y axis) and number of clusters (black line, left Y axis) as an indicator for the self-sorting process over time. While the number of total Sox17 cells keeps rising, the number of clusters starts to decrease due to merging of cell clusters. Clusters were automatically determined as isolated surfaces (see Methods) (F) FACS analysis of Sox17-RFP Bra-GFP EBs at 48 (left) or at 72 (middle) and 120 (right) hours of differentiation. (G) A model of the dynamics observed for Sox17 onset and expansion.

It was previously shown that only Brachyury+ cells give rise to Sox17+ cells in differentiating mESCs [22]. We first verified that Sox17 cells uniquely originate from Bra+ cells in our system. We analyzed by FACS the composition of EBs at 48, 72 and 120 hrs after induction by CHIR. After the emergence of a GFP-only population (48hrs), a double-positive population emerges (72 hrs), followed by an RFP-only population (120 hrs) (Fig. 3f). We plated GFP+/RFP-, GFP+/RFP+ and GFP-/RFP+ populations sorted at the 72 hrs time point, and imaged them 24 hours after the sorting. The GFP+/RFP-population showed no fluorescent signal, and displayed a mesenchymal morphology, suggesting a later mesodermal fate. In contrast, both the GFP+/RFP+ and GFP-/RFP+ populations showed only an RFP signal and a rounded morphology (Fig. S5A). FACS sorting of the cells from the 2D differentiation protocol led to similar results (Fig. S5B). Together, these results suggest that Sox17 is activated within a subset of the Bra-GFP+ cells in the studied in-vitro models, followed by self-sorting dynamics of these cells (Fig. 3g).

### Sox17+ cells self-sorting is correlated with E-cadherin expression

The differentiation of mesendoderm cells to DE is known to be associated with re-activation of E-cadherin [23, 24]. We hypothesized that the self-sorting movements of the Sox17-RFP+ cells are correlated with the mesenchymal to epithelial transition (MET) and E-Cadherin (E-cad) expression. It has been shown on mutant mice that Sox17 is required for DE cell egression, and that E-cad was differentially expressed in egressing DE cells compared to cells that fail to egress [10, 25]. In our systems, Sox17+ cells that have already migrated to the outer layer of the EB or have formed a confined lumen show the highest E-cad expression (Fig. 4a,b). Sox17+ unsorted cells show low to no E-cad expression (Fig. 4b,c, Movie S4). Outer layer EB cells that are Sox17-, do not express E-cad (Fig. 4d). Moreover, at a later stage, when all Sox17 cells are sorted, the high E-cad expression is seen in all Sox17+ cells and is limited to Sox17+ cells only (Fig. 4e,f Movie S5).

**Figure 4.**
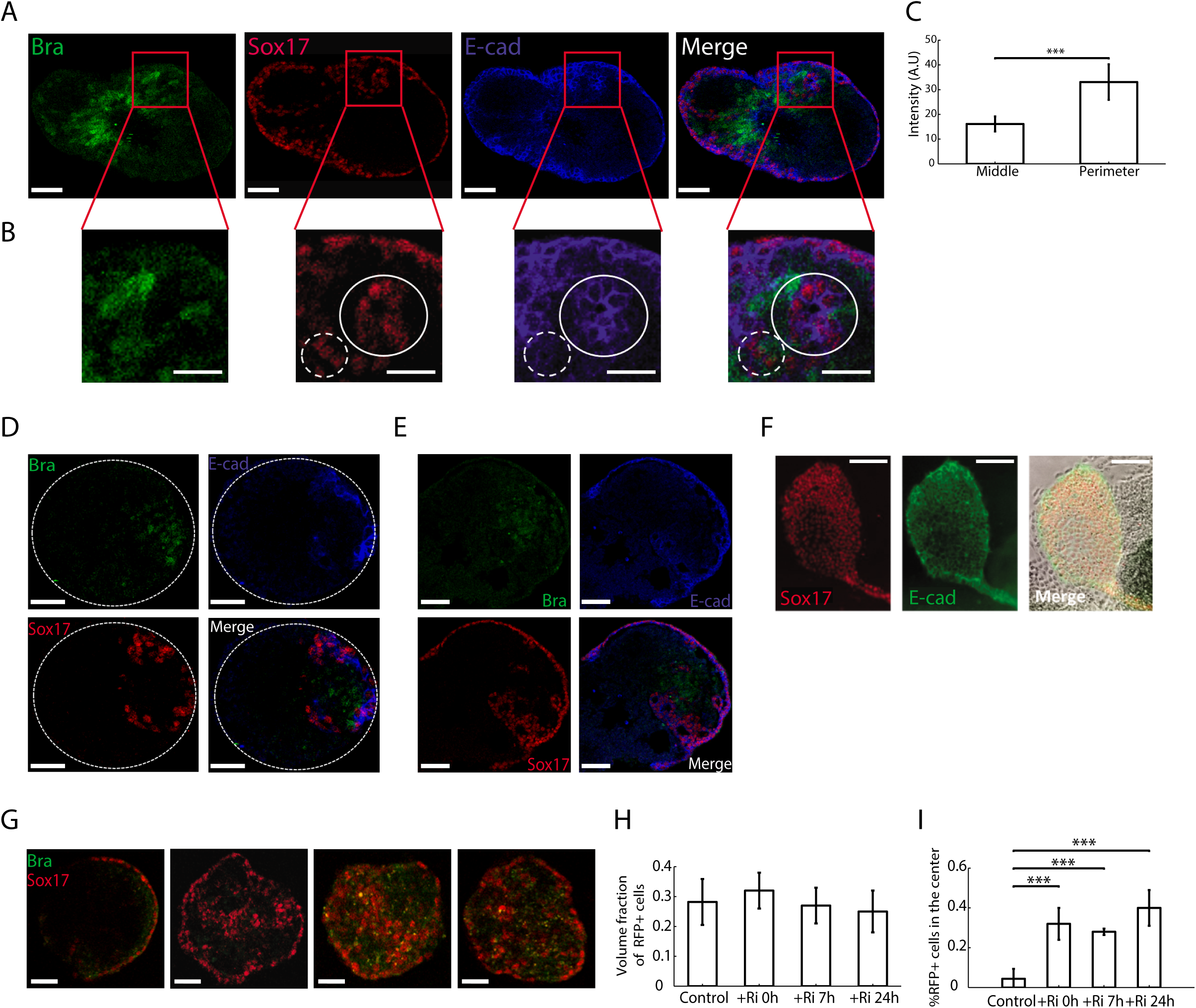
Self-sorting is not required for establishment of Sox17+ population, and is concomitant with E-cad upregulation. (A) E-cad immunostaining at 48 hours of differentiation. E-cad is restricted to the Sox17 cells and not the Bra expressing cells. The highest E-cad expression is obtained in the Sox17 cells surrounding the outer layer of the EB or forming an interior lumen. Scale bar 50um. (B) Zoom-in in the same representative EB. E-cad expression level depends on Sox17 cells location and aggregation stage. Scale bar 25um. (C) E-cad intensity in the cells in the middle of the EB is significantly lower that on the perimeter of the EB (n=10, P = 10^-5^) (D) E-cad is restricted to the Sox17 cells on the outer layer of the EB. Scale bar 50um. (E,F) E-cad expression of Sox17 cells in 72 hours differentiated EBs (E) or 2D colonies (F). Scale bar 50um and 100um respectively.(G) Representative EBs at 65 hours into differentiation, without (left) or with addition of ROCK inhibitor (Y-27632, 20uM) at 0hrs, 7hrs or 24hrs after the addition of CHIR (Left to right). (H) Number of Sox17 cells per EB, normalized to the EB volume, for different timings of ROCK inhibitor addition to the medium. (I) Fraction of Sox17+ cells in the middle of the EB out of all Sox17 cells, for different ROCK inhibitor addition timings. (n=8). *** P < 0.001 ** P < 0.01 * P < 0.05

These results suggest that Sox17+ cells dispersed in the EB start to accumulate E-cad, and possibly the low concentration of E-cad expression is sufficient to initiate self-sorting. The aggregation of the Sox17+ cells results in an upregulation and higher expression of E-cad. To test the effect of cell movement and resulting aggregation on the final distribution of Sox17+ vs. Sox17-populations, we blocked cell movement by adding the Rho-associated protein kinase inhibitor Y-27632 (ROCKi) at different time points during differentiation. As expected, the addition of ROCKi reduced significantly the self-sorting of the Sox17+ cells, resulting in a higher fraction of cells in the interior of the EB at 65h compared to the control, where these interior cells are not forming lumens (Fig. 4h,i). The final fraction of Sox17+ cells in each EB was not altered by the addition of ROCKi (Fig. 4g). Taken together, these results show that self-sorting and increase in the expression of E-cad occur after cells already express Sox17, and that this self-sorting and aggregation is not required for Sox17 onset and proliferation. Following this conclusion, we next wanted to evaluate the dependence of Sox17 onset on signaling pathways.

### Temporal requirements of Wnt and Nodal/Activin signaling for Sox17 specification

In the embryo, Activin/Nodal signaling is required for endoderm differentiation [22, 26, 27]. Canonical Wnt TCF/beta-catenin were also shown to directly bind to the Sox17 promoter during endoderm differentiation [28]. In our system, though external inducing signals (e.g. CHIR) are supplied as a uniform field, cells still respond variably, either due to differential activation and response of signaling pathways or alternative chromatin states. To explore the interplay and temporal role of the Wnt and Nodal signaling pathways in DE specification under these conditions we tested the effect of various signal perturbations on the differentiation process. As it was shown that a 24-hours pulse of Wnt activation is sufficient in gastruloids for the induction of both Bra and Sox17, we first tested the needed duration of Wnt induction in our system [14, 15]. Induction with CHIR for 24 hours, or even 12 hours, is sufficient for the expression of both Bra and Sox17 (Fig. 5a top row; Fig. S6A top). Replacing CHIR at 24 hours with an inhibitor of endogenous Wnt signaling (using the porcupine inhibitor IWP2, thus inhibiting Wnt ligand secretion) reduced the Sox17+ colonies in a GFP intensity dependent manner (Fig. 5a middle rows,b,c,d). This suggests that some of the GFP-high cells (corresponding to longer or stronger activation of Brachyury) are already committed to their Sox17 fate regardless of the Wnt pathway inhibition. This effect is abolished when IWP2 is added 12 hours after CHIR (Fig. S6). When IWP2 is added after 24 hours while maintaining CHIR, all the Bra+ colonies give rise to Sox17 populations, while Bra-colonies remain negative (Fig. 5a bottom row,b,c). Taken together these results suggest that a sub population of Bra+ cells gain their potential to differentiate to a Sox17+ state shortly after Bra onset. Cells that are already committed to their differentiation do not depend on the activation of Wnt signaling anymore, and can differentiate under its inhibition at this point. Bra+ cells that are not committed yet to activate Sox17 will fail to further differentiate to a Sox17+ state.

**Figure 5.**
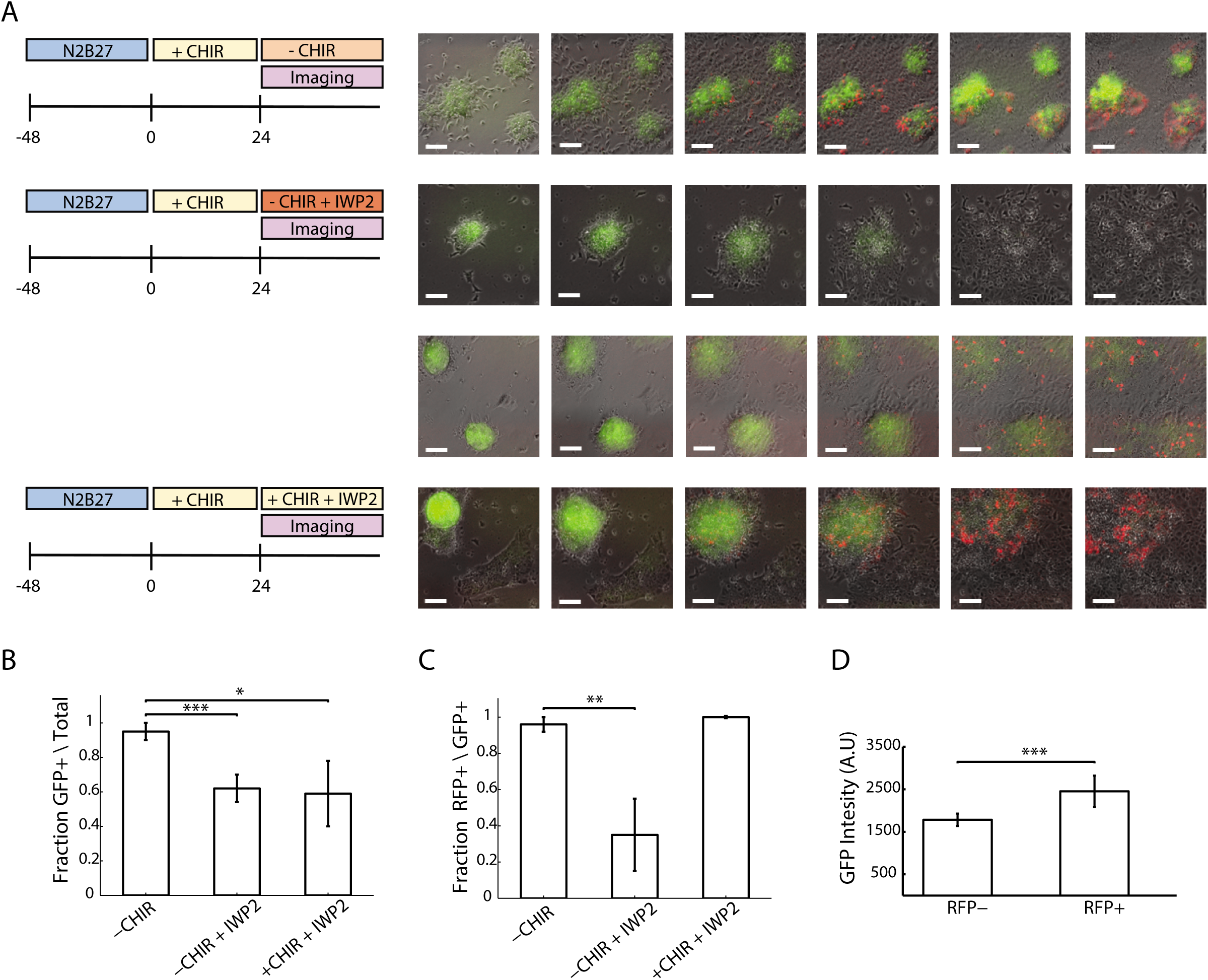
Sox17+ establishment is independent of Wnt in Bra-high cells. (A) Time lapse images of Bra and Sox17 expression over time under withdrawal of CHIR at 24 hrs (top), withdrawal of CHIR and blocking the Wnt pathway (IWP2) (middle) or blocking the Wnt pathway (IWP2) with CHIR in the medium (bottom). (B) Fraction of colonies with GFP+ cells out of all the colonies for each condition. (C) Fraction of colonies with RFP+ cells out of all colonies with GFP+ cells. (D) GFP intensity at 24 hours, in RFP+ and RFP-colonies (at 60 hours).

It was previously shown that treatment with Activin allows DE differentiation and Sox17 expression [14, 29]. In our system, the Wnt signaling activation as an external field is sufficient for Sox17 emergence. We therefore wanted to dissect the role of Wnt signaling and Activin\Nodal signaling on the onset of Sox17. Using a different double reporter cell line with Wnt (SuTOP-CFP) and Nodal/Activin (AR8-RFP) pathway reporters, under our standard differentiation protocol (Fig. 2a) we find that CHIR allows activation of the Activin\Nodal pathway in our in-vitro system (Fig. S7A). Interestingly, the Wnt activity concentrated in the middle of the colony while the Activin reporter cells are sparsely distributed within and at the edges of the colony, resembling the expression pattern of Bra and Sox17, respectively (Fig. S7A). Indeed, the majority (70+-9.9%) of Sox17+ cells are co-localized with or immediately adjacent to AR8-RFP+ cells, suggesting a differential activation of the Activin\Nodal signaling pathway within the Bra+ population can partially explain the Sox17 onset pattern we see (Fig. S7B,C). When replacing CHIR with Activin as an external signal the onset timing of the Sox17 cells was delayed (Fig. S7D). When replacing CHIR at 24 hours with the Activin\Nodal pathway inhibitor SB431542 (SB), the potential of Bra+ colonies to differentiate to Sox17+ cells was reduced by ∼3-fold and was not rescued by the addition of CHIR (Fig. S7E,F). Adding SB with CHIR from the beginning results in no mesodermal nor endodermal differentiation (Fig. S7G), similar to what was shown *in vivo* with a Nodal null mutant [26]. These results suggest that both Wnt and Activin signaling pathways are needed for the initial differentiation of Bra, and commitment for DE emergence. While Wnt signaling pathway is crucial for Bra differentiation and Wnt signaling inhibition at mature Bra cells allows Sox17 expression, Activin\Nodal is required for the expression of Sox17 throughout the differentiation process, with correlated spatial patterns between the two.

## Discussion

Here we explored the onset and expansion dynamics of mesendoderm-derived DE cells. We show that in the embryo, the first Sox17-RFP+ cells arise in a dispersed pattern, though limited to the anterior PS, followed by expansion and migration. In both 2D and 3D in-vitro cultures, we then demonstrated that Sox17+ cells emerge in a stochastic salt-and-pepper pattern within the Bra+ cell population. In this sense, the in-vitro conditions expand the Sox17-competent region compared to the embryo, while maintaining the disperse pattern. This expansion may be attributed to more uniform signals sensed by the cells in the in-vitro systems, compared to the embryo, where localized sources of Wnt and Nodal signaling result in spatial gradients of these signals. This onset pattern is in sharp contrast to the strongly-localized pattern of Brachyury onset [14, 17, 30]. The ordered pattern of Sox17+ cells is then obtained through self-sorting of the cells. The transition from stochastic to self-sorted pattern is emphasized in the in-vitro systems due to the spatial expansion of the Sox17-competent region. These results suggest that in the embryo, mesendoderm progenitors, while egressing through the primitive streak, stochastically commit to definitive endoderm, and then through cadherin-mediated self-sorting, get spatially sorted apart from mesoderm cells [10, 31]. In the embryo, the sparse DE cells egress at this stage and intercalate into the VE layer, while in the in-vitro systems, in the absence of VE, they cluster together to form lumens or an envelope layer. In our in-vitro models, the signaling gradient patterns and signal accessibility in different regions differ greatly between the 2D and 3D setups [32]. The robustness of the relative temporal pattern between Bra and Sox17 and the spatial stochasticity in DE onset in these two model types therefore suggest it is internal state maturation that determines which cells will adopt a DE fate, as opposed to differences in external signaling.

We find that E-cadherin expression, or at least the amount of membrane-bound E-cadherin, increases in parallel to the self-sorting of Sox17+ cells. This is consistent with previous observations in the embryo, where high E-cadherin is observed in DE cells only after they have egressed and intercalated into the VE [10, 25]. The sparse, spatially stochastic appearance of DE cells, along with the late induction of E-cadherin may facilitate the intercalation process, preventing pre-mature formation of DE cell clusters or lumens within the epiblast, as they are formed in the EBs. Once self-sorting is completed in the EBs or 2D colonies, E-cadherin expression is unique to the Sox17+ populations. Limiting cell movement, by inhibition of Rho-associated protein kinase (ROCK), an important controller of actin microfilaments, did not affect the differentiation to Sox17+ cells nor their proliferation, regardless of the timing of inhibition. It was previously shown that ROCK inhibition and cytoskeletal reorganization can mediate activation of Wnt signaling and the induction of EMT, and can affect cell differentiation in vitro [33–35]. While ROCK inhibition early in differentiation may induce EMT and mesendoderm differentiation and by that enhance Sox17 expression later on, the emergence of a similar fraction of Sox17+ cells when ROCKi is added later in differentiation indicates that Sox17 expression does not require cell movement. The order of Sox17 onset, E-cadherin upregulation and cell movement suggests the activation of Sox17 does not depend on the latter two, but may drive these processes.

In the Embryo, proximal-distal Nodal gradient and posterior-anterior Wnt gradient allow pattern formation during gastrulation [5]. In vitro, it was shown that a 24 hours pulse of CHIR (with or without Activin) induces EB gastrulation like events, including Bra and Sox17 expression [14]. Moreover, it was shown that Activin induces DE differentiation and Sox17 expression in Bra+ cells [22, 36]. Here we showed that even a short Wnt activation is sufficient to induce Bra, and shortly after, a small sub-population of the Bra+ cells is already committed to their Sox17+ fate independently of Wnt activation. Our results further suggest that a disperse local activation of the Activin\Nodal pathway may partially explain this altered subpopulation, but cell-to-cell variation in expression or chromatin state may also account for the differential DE potential. A better understanding of the specific chromatin and expression determinants of this potential will provide a comprehensive picture of this spatially-stochastic cell fate decision. Nonetheless, our *in vivo* and *in vitro* results highlight basic rules governing DE onset and patterning through the commonalities and differences between these systems.

## Acknowledgments

We thank Z.D. Smith and A. Arczewska for help with CRISPR knock-in design and implementation. This work was supported by the Israeli Science Foundation (1665/16), the New York Stem Cell Foundation, the NIH (P01 GM099117 and P50 HG006193) as well as the Max Planck Society.

## Author Contributions

MP and IN designed and conceived the study. MP and ASK performed experiments. LW and MW assisted with tetraploid aggregation and embryo isolation. MP performed the data analysis. IN and AM oversaw the project. MP and IN wrote the paper with contributions from ASK and AM.

## Declaration of Interests

Nothing to declare.

## Methods

### Cell culture

Mouse ESCs were cultured on gelatin surface using standard conditions on irradiated primary mouse embryonic fibroblasts and knockout DMEM containing 15% fetal bovine serum, 50 ug/ml penicillin/streptomycin, 2 mM L Glutamine, 100 μM non-essential amino acids, 80 μM ß-mercaptoethanol and 10^3 U/mL LIF. Cell lines used are E14 Bra-GFP mouse ESCs (kindly provided by Dr. Gordon Keller), AR8-RFP\SuTOP-CFP mouse ESCs (kindly provided by Dr. Palle Serup).

### Sox17-RFP live marker

A sox17 homology donor (SOX17-HomDon) plasmid was generated, containing (i) a left homology arm containing a 1-kb sequence immediately upstream of the SOX17 stop codon; (ii) a P2A-H2B-Strawbwry cassette; (iii) a floxed Hygromycin selection cassette (loxP-PGK-Hygro-loxP); and (iv) a right homology arm containing a 1-kb sequence immediately downstream of the SOX17stop codon. A single-guide RNA (sgRNA) recognizing a sequence near the stop codon of SOX17 (‘GCAACTACCCCGACATTTGA’) was cloned into a Cas9-nickase expression vector pSpCas9(BB)-2A-Puro (PX459) from the Zhang laboratory, Addgene plasmid # 62988). This plasmid, together with the SOX17-HomDon plasmid, was transfected into E14 Bra-GFP cells using Xfect™ transfection reagent (Cat. # 631318, Takara). After selection, single clone was validated with sequencing from the genomic Sox17 locus. After Cre-floxing out of the Hygromycin selection marker, a single clone was validated again, expended and frozen in aliquots for the use in all experiments.

### Differentiation essay

Cells were cultured in serum-free N2B27 media supplemented with LIF and and 2i (3 μM CHIR99021 and 1 μM PD0325901) for a minimum of 24 hours. Prior to 2D differentiation, ESCs were fully dissociated to single cells and seeded at low density (∼12,000 cells/cm2) on gelatin-coated plates in LIF 2i medium supplemented with 5% KSR. Differentiation was induced a minimum of 18h later by rinsing ESCs thoroughly with 1X PBS and changing to N2B27 media without LIF 2i. For 3D Embryoid bodies’ aggregation, single cells were transferred to a low-adherent culture dish, allowing cells to randomly aggregate together in small clumps, in N2B27 media without LIF 2i. In both 2D and 3D differentiation protocols, after 48hrs in in N2B27, media was changed to N2B27 media with CHIR99021 (3 μM).

For Wnt\Activin signaling pathways interplay experiments, cells were supplemented with CHIR99021 (3 μM) or IWP2 porcupine inhibitor (10μM) or Activin (20ng/ml) or SB-431542 Activin receptor inhibitor (10μM), as indicated in the relevant result.

### Live imaging

2D differentiation experiments were imaged using a Nikon TiE epi-fluorescence microscope equipped with a motorized XY stage (Prior) and taken within a connected 6 × 6 or 7 × 7 spatial range at 10× magnification in up to three fluorescent wavelengths and phase contrast using NIS Elements software.

3D differentiation experiments were imaged using a Zeiss LSM7 inverted two-photon microscope with a 20X/0.8NA air objective. Each EB was scanned at 3um intervals along the *z* direction. Horizontal resolution was set to 512×512 pixels at approximately 0.6um per pixel. GFP was excited at 920nm. RFP was excited at 760nm, Alexa 488, Alexa 405 and Alexa 594 were excited at 800nm. CFP was excited at 860 nm.

For both systems, acquisitions were taken every 30-60 minutes for 2-5 days. For live imaging, an Okolab incubation cage, maintaining 5% CO2 and 37C was used.

### Immunohistochemistry

For immunostaining, cells were fixed in 4% paraformaldehyde and immunostained for the indicated primary antibody at the concentration recommended by the manufacturer, at 4C O\N (unless indicated otherwise). For 2D colonies, cells were then washed 3 times in PBS and incubated with a secondary antibody for one hour at RT. For 3D EBs were washed in four cycles in blocking buffer (PBS, 0.15% Triton x100, 7% FBS) for at least half an hour per cycle. A secondary antibody in blocking buffer was incubated o/n, followed by four wash cycle as described for the primary antibody. Antibodies used: Sox17 - R&D Systems AF1924 (3 h at RT); Bra - R&D Systems AF2085; E-cadherin - Cell Signaling 24E10.

### Aggregation of mouse embryonic stem cells and Animal procedures

Tetraploid complementation was performed with mESCs thawed from a frozen vial and cultured for three days on a layer of CD-1 feeders (Artus, J. & Hadjantonakis, A. K.). Aggregated embryos were re-transferred bi-laterally in a clutch of 10-15 embryos into uterine horn of CD-1 strain pseudopregnant foster females. All animal procedures were performed in accordance with the institutional, governmental and state regulations (LAGeSo Berlin, G0243/18).

### Embryo isolation and imaging

Embryos from the uterine decidua were dissected in cold 1X HBSS and the Reichert’s membrane was carefully removed with a thin-drawn glass capillary. The embryos were transferred through three washes in 1X PBS with 0.4% BSA and fixed overnight in 4% PFA at 4°C. The embryos were washed three times of 5 minutes each in 1X cold PBS with 0.1% TritonX-100. Nuclei were counterstained with 0.24μg/mL of DAPI in 1X PBS with 0.25% Triton X-100 at 4°C for 2 hours, followed by three washes with 1X PBS with 0.1% TritonX-100. Images were acquired with a Zeiss LSM880 AiryScan confocal microscope at 10X magnification with a z-stack of 2.73μm interval and averaging of 8 frames. Images were processed Imaris and Fiji.

### Quantitative Real-Time PCR

The cells were washed with cold 1xPBS and lysed with 300 - 500μL of TRIzol reagent. Equal volume of 100% ethanol was added to the lysate, mixed by vortexing and passed through a RNA-bind column. DNaseI treatment for 15mins was performed directly on the column to remove genomic DNA contamination. RNA was isolated as per manufacturer’s instructions (RNeasy Mini Kit). Reverse transcription was performed as per manufacturer’s instructions (RevertAid first strand cDNA synthesis kit) with 1μg of RNA. Quantitative real-time PCR was performed with intron-spanning primers for *T* (GGTCTCGGGAAAGCAGTGGC; CATGTACTCTTTCTTGCTGG) and *Sox17* (GGAGGGTCACCACTGCTTTA; AGATGTCTGGAGGTGCTGCT).

### Flow cytometry

Flow cytometry for differentiation kinetics of the reporters was performed on the BD Accuri C6 flow cytometer. 100,000 events were acquired and the cells were gated for size and singlets using forward (FSC-A/W) and side scatter (SSC-A/W). GFP and RFP was measured by excitation of blue and yellow-green laser and filtered through 530/30 and 610/20 band pass filter respectively. The gated data was exported and processed using FlowJo version 10.

### Segmentation and Image Analysis

The Two-photon microscopy data was contrast enhanced in Fiji and then spot segmented in Imaris, where the integration radius was set to 7um and the filter was set to sum intensity with a threshold of 110/255. The background is subtracted prior to segmentation. The segmented data was read and analyzed in MATLAB. Sox17 onset were set when 7 segmented RFP+ cells above intensity threshold were detected. For self-sorting analysis of the Sox17-RFP+ cells, ‘Surface’ Imaris tool were used.

### Statistical analysis

For the analysis of Sox17-RFP+ cells onset pattern, segmented RFP cells and segmented GFP cells were analyzed for their location at Sox17 onset time point within the EBs. Images were cleaned from detected clumps of dead cells outside of the EB radius in order to prevent misleading distance measurements. The distances between every two RFP+ cells as well as between every two GFP+ cells were then calculated. The distributions of the distances were compared using two-sample Kolmogorov-Smirnov test under the null hypothesis that the two sets of distances are from the same continuous distribution. The results represent the analysis of 5 EBs of similar size.

**Figure S1.**
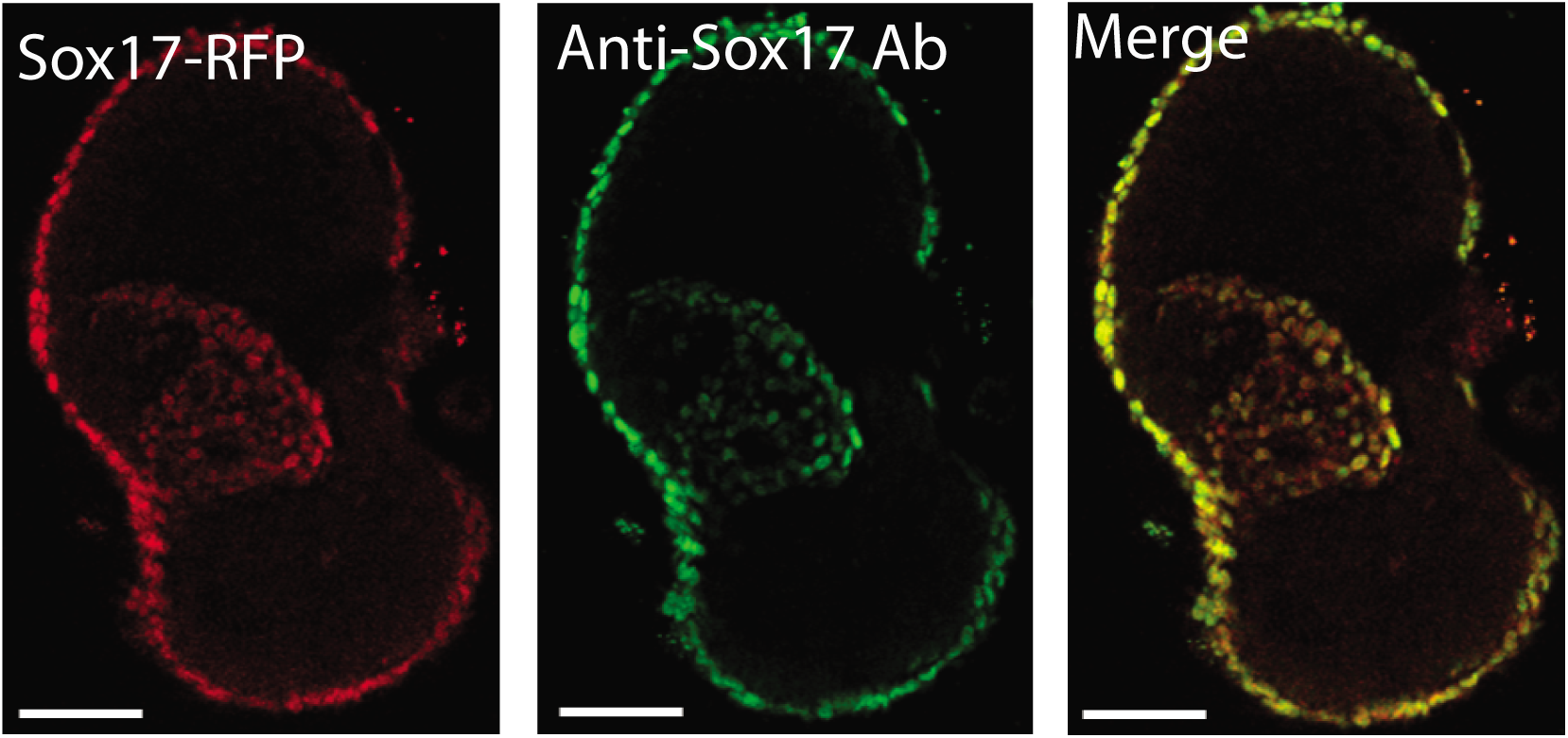
Verification of the Sox17 live marker in EBs. Overlap between Sox17-RFP live marker (left) and Sox17 antibody immunostaining (middle) to validate the live marker cell line in a 3D EB. Scale bar 50um.

**Figure S2.**
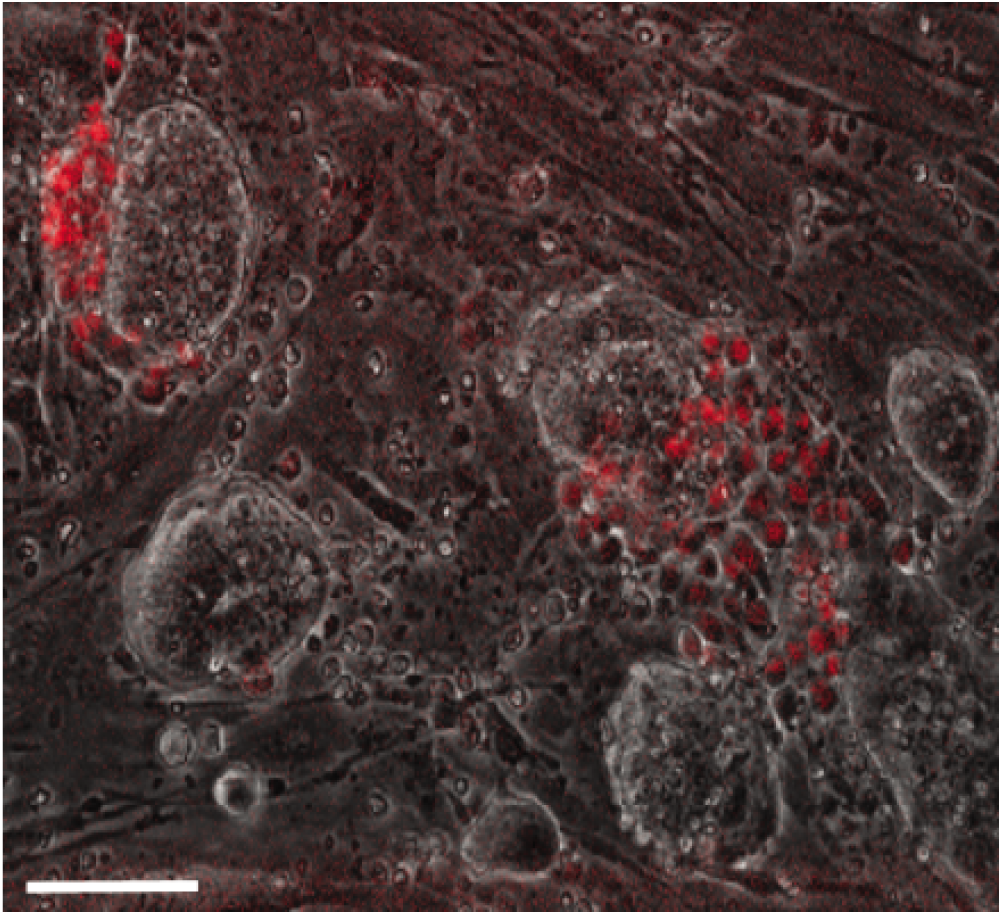
Sox17 peripheral expression in 2D pluripotent colonies. Sox17-RFP cells in ES 2D colonies under pluripotency conditions. Scale bar 100um.

**Figure S3.**
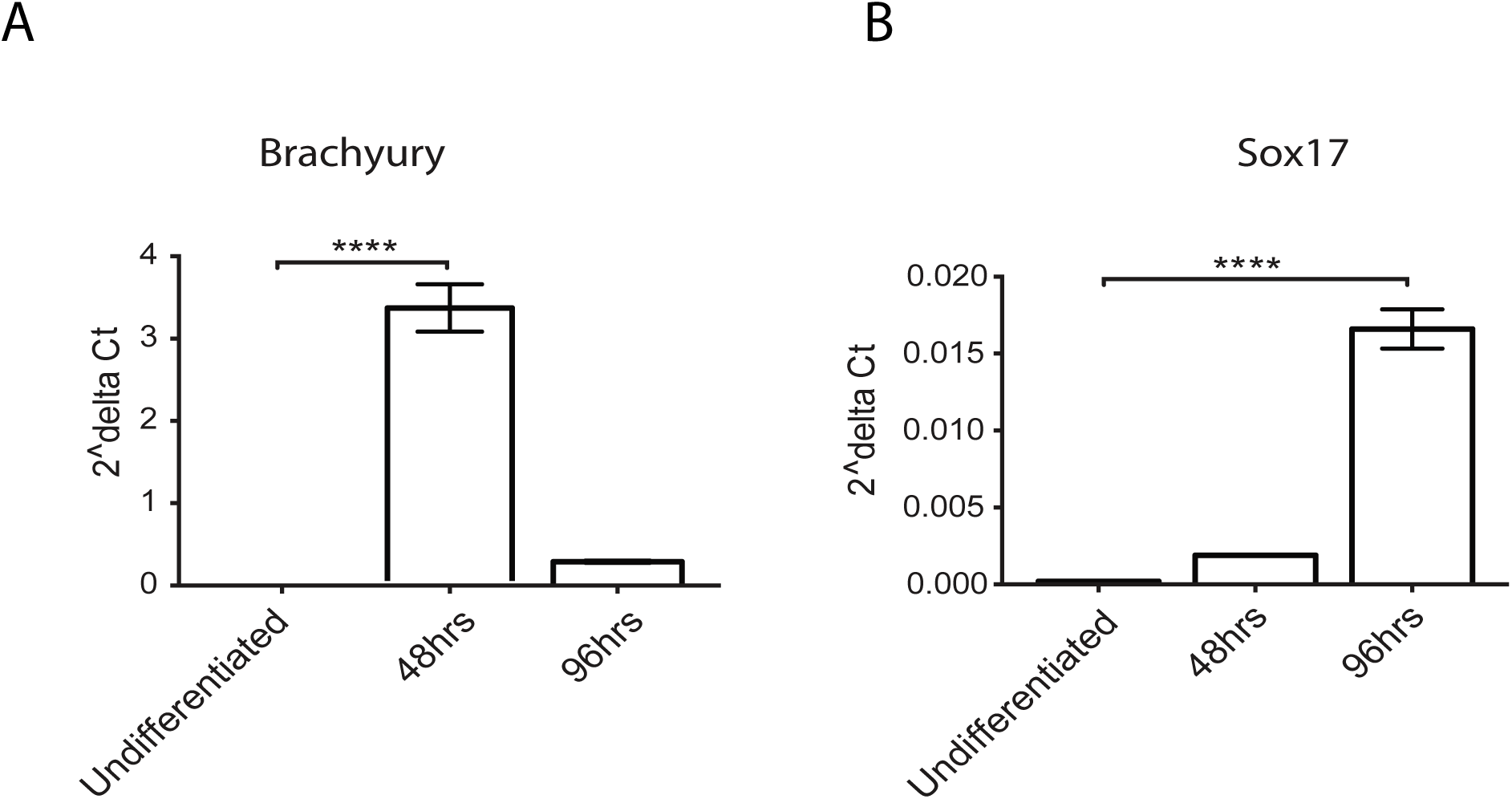
Bra expression precedes Sox17 at the RNA level. (A,B) RT-qPCR results for expression of Bra (A) and Sox17 (B) at undifferentiated, 48 hours or 96 hours of EB differentiation, show a time interval between Bra and Sox17 at the RNA level.

**Figure S4.**
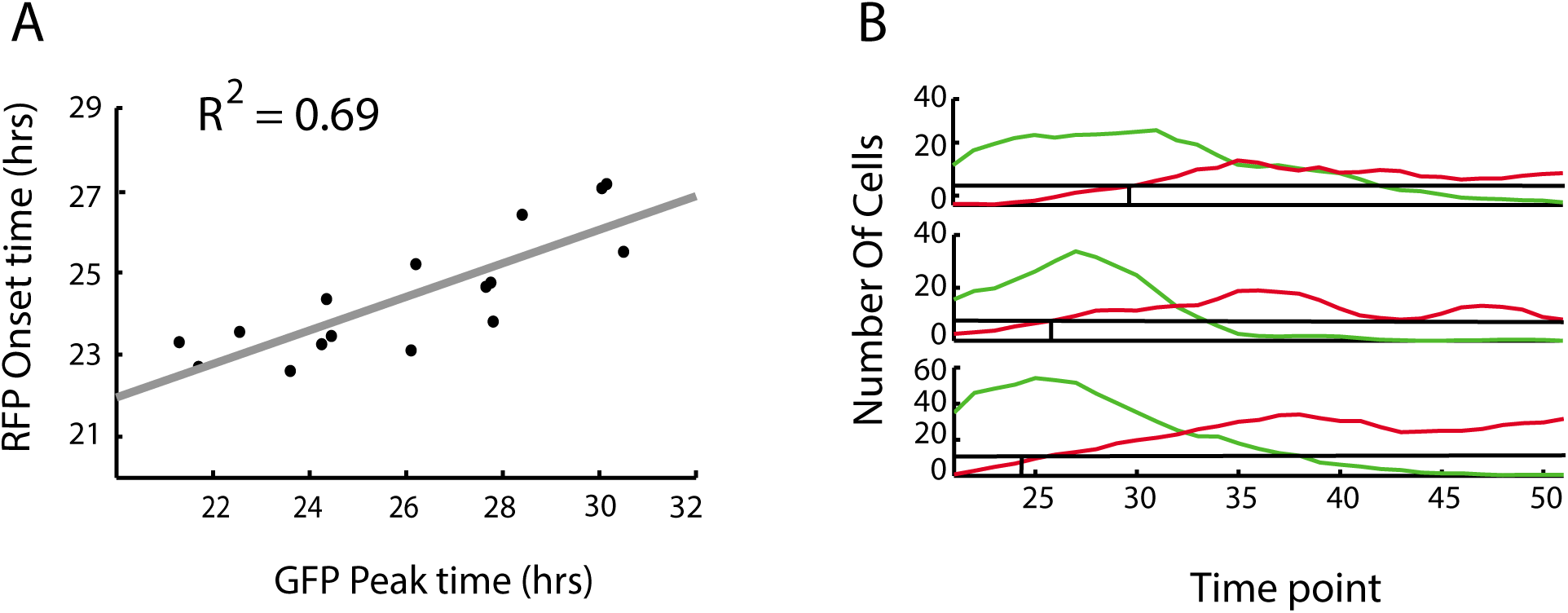
A regular temporal gap between Bra peak and Sox17 onset. (A) Bra-GFP Peak time versus Sox17-RFP Onset time, each dot represents one EB (n=18). (B) Number of Bra-GFP (green) cells or number of Sox17-RFP (red) cells per EB over time in 3 EBs as an example for the coupling between Bra peak and Sox17 onset. Later GFP peak time show later RFP onset time.

**Figure S5.**
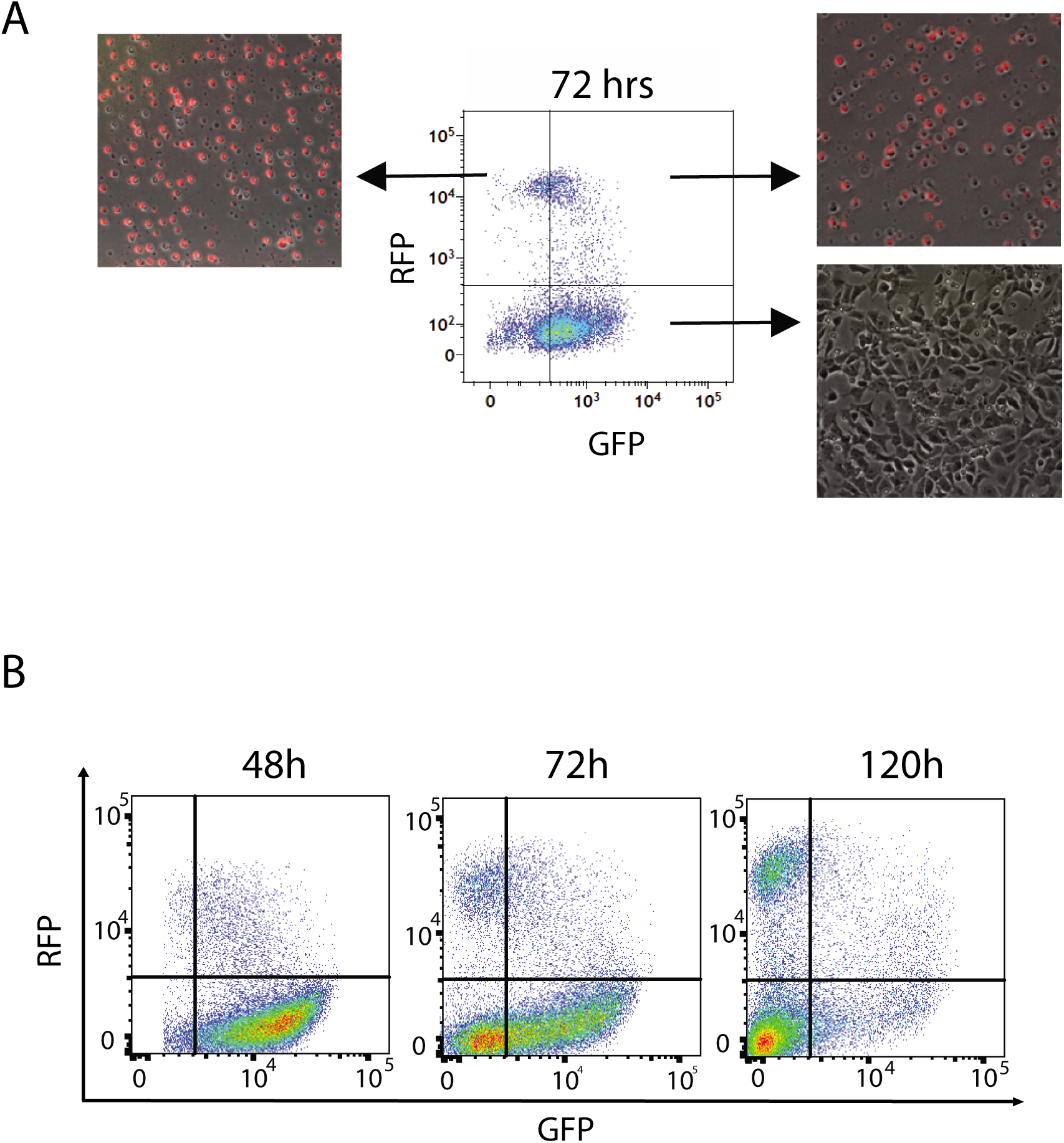
Cells differentiated for 72 hours show two distinct populations. (A) EBs differentiated for 72 hours were FACS sorted to 3 populations (GFP+, RFP+ and GFP+ RFP+), plated and imaged after 24 hours. GFP only cells (right, bottom) show a mesenchymal morphology and no GFP signal. Double positive and RFP only cells (Top, right and left, respectively) show a similar round morphology with low attachment properties, and RFP signal in most cells. (B) FACS analysis of Sox17-RFP Bra-GFP 2D colonies at 48 (left) 72 (middle) and 120 (right) hours of differentiation.

**Figure S6:**
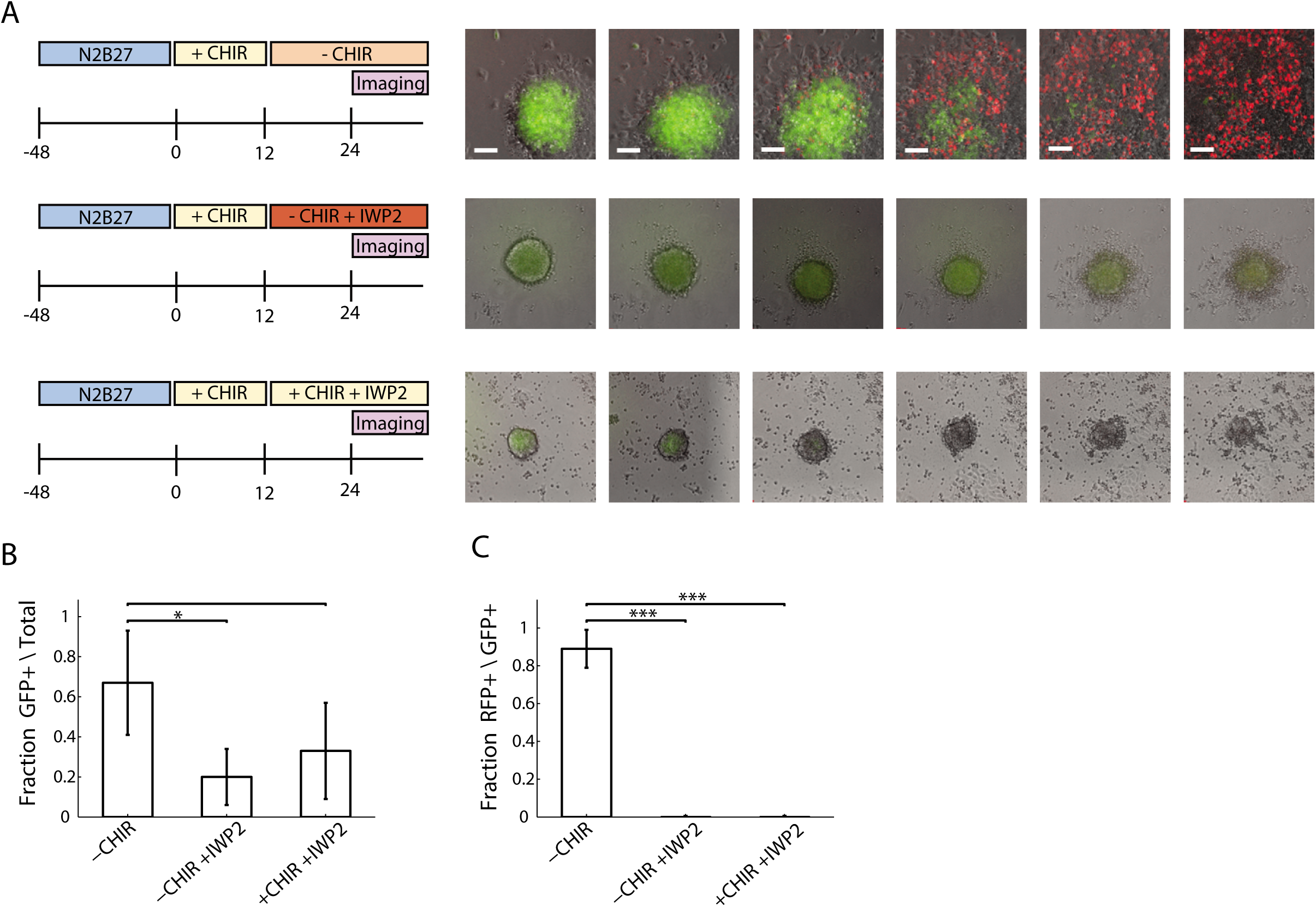
Wnt signaling pathway inhibition at 12 hours of differentiation. (A) Time lapse images of Bra and Sox17 expression over time under withdrawal of CHIR after 12 hrs (top), withdrawal of CHIR and blocking the Wnt pathway (IWP2) (middle) or blocking the Wnt pathway (IWP2) with CHIR in the medium (bottom). (B) Fraction of colonies with GFP+ cells out of all the colonies for each condition. (C) Fraction of colonies with RFP+ cells out of all colonies with GFP+ cells.

**Figure S7:**
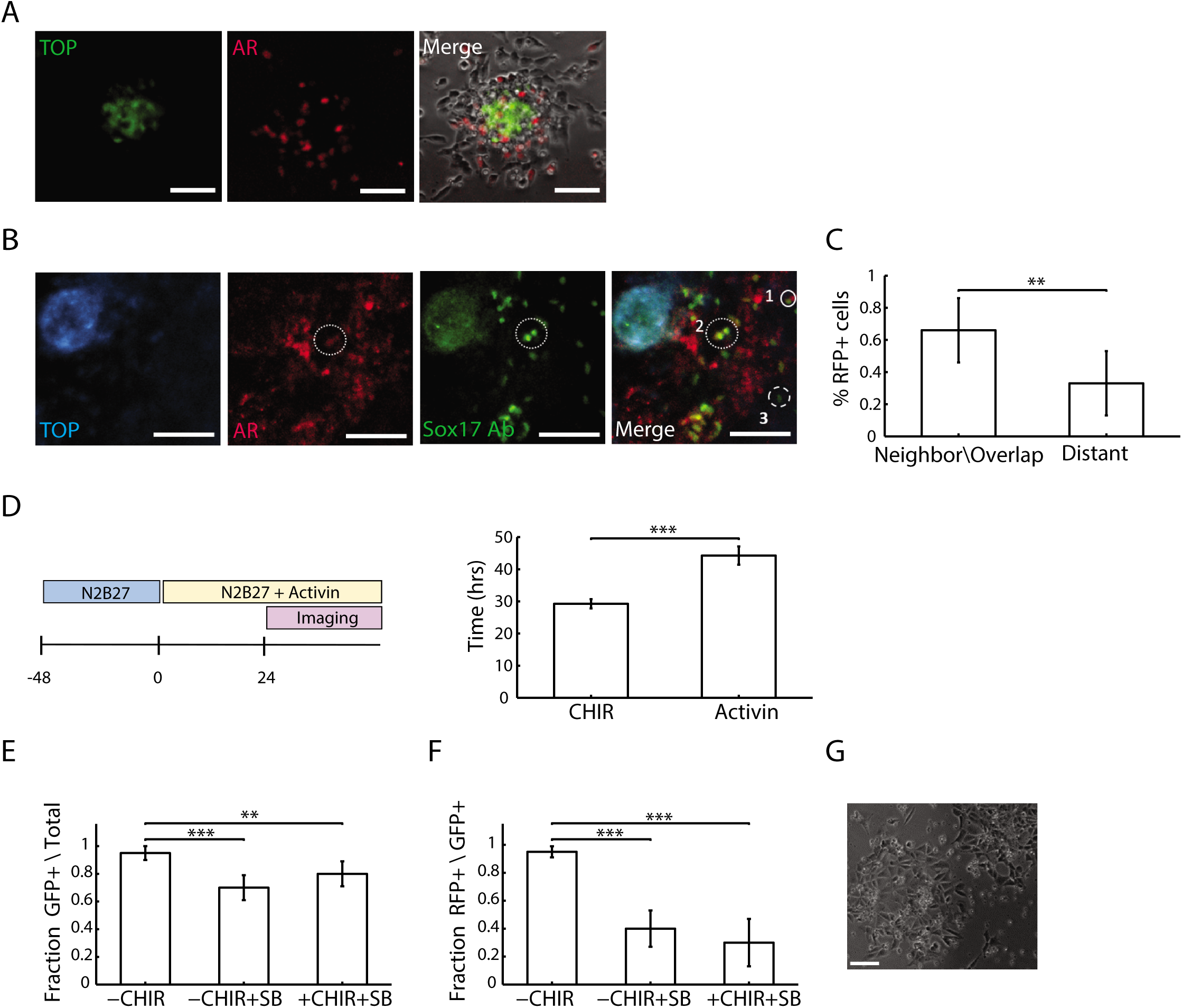
The interplay between Wnt and Nodal\Activin signaling pathways in the specification of Bra cells into Sox17 cells. (A) AR-RFP (Activin signaling reporter, red)-TOP-CFP (Wnt signaling reporter, green) cells were imaged at 48 hours of differentiation. Scale bar 100um (B) Sox17 expressing cells (immunostaining, green) are nearest neighbor to (1), overlap (2), or distant from (3) AR+ cells. Scale bar 50um (C) Fraction of Sox17+ cells in a colony that exhibit different relations to AR+ cells (average and SD over n=5 colonies). (D) Timing of RFP onset at +CHIR or +ACT conditions. (E,F) Fraction of colonies with GFP+ cells from total colonies (E) or RFP+ cells from colonies with GFP+ cells (F) at withdrawal of CHIR after 24 hours, withdrawal of CHIR and blocking the Nodal\Activin pathway (SB431542, 10uM) or blocking the Nodal\Activin pathway with CHIR in the medium. (G) Addition of SB with CHIR at the beginning of the experiment results in no Bra or Sox17 expression. Scale bar 50um.

**Movie S1. Dynamics of Sox17-RFP onset and expansion in an embryoid body.**

Three-dimensional time-lapse imaging of Bra-GFP, Sox17-RFP in an embryoid bodies, imaged between 24 and 72 hrs of differentiation. Left: top view of raw signal. Middle: RFP and GFP segmentation. Right: surface-rendering for the RFP signal. Scale bar 50 um.

**Movie S2. Dynamics of Sox17-RFP onset and expansion in 2D colonies.**

Three colonies imaged between 24 and 57 hrs from transition to differentiation medium.

**Movie S3. Aggregation dynamics of Sox17-RFP+ cells in EBs.**

Dynamics of Sox17 onset and clustering in five EBs. Left: top view of raw signal. Right: surface-rendering for the RFP signal. Scale bar 50 um.

**Movie S4. E-cadherin expression concomitant with cell movement.**

Z-stack (2um) image of Bra-GFP, Sox17-RFP in an embryoid body, immunostained for E-cad (blue) at 48 hours after CHIR induction. Scale bar 100 um.

**Movie S5. E-cadherin expression is limited to all sorted Sox17 expressing cells.**

Z-stack (2um) image of Bra-GFP, Sox17-RFP in embryoid body, immunostained for E-cad (blue) at 72 hours after CHIR induction. Scale bar 100 um.

## References

1. Tam PP, Loebel DA (2007) Gene function in mouse embryogenesis: get set for gastrulation. Nat Rev Genet 8: 368–381

2. Singh AM, Reynolds D, Cliff T, Ohtsuka S, Mattheyses AL, Sun Y, Menendez L, Kulik M, Dalton S (2012) Signaling network crosstalk in human pluripotent cells: a Smad2/3-regulated switch that controls the balance between self-renewal and differentiation. Cell Stem Cell 10: 312–326

3. Yoney A et al (2018) WNT signaling memory is required for ACTIVIN to function as a morphogen in human gastruloids. Elife 7:

4. Robertson EJ (2014) Dose-dependent Nodal/Smad signals pattern the early mouse embryo. Semin Cell Dev Biol 32: 73–79

5. Norris DP, Robertson EJ (1999) Asymmetric and node-specific nodal expression patterns are controlled by two distinct cis-acting regulatory elements. Genes Dev 13: 1575–1588

6. Shen L et al (2007) Genome-wide profiling of DNA methylation reveals a class of normally methylated CpG island promoters. PLoS Genet 3: 2023–2036

7. Yamamoto M, Saijoh Y, Perea-Gomez A, Shawlot W, Behringer RR, Ang SL, Hamada H, Meno C (2004) Nodal antagonists regulate formation of the anteroposterior axis of the mouse embryo. Nature 428: 387–392

8. Niakan KK et al (2010) Sox17 promotes differentiation in mouse embryonic stem cells by directly regulating extraembryonic gene expression and indirectly antagonizing self-renewal. Genes Dev 24: 312–326

9. Kanai-Azuma M et al (2002) Depletion of definitive gut endoderm in Sox17-null mutant mice. Development 129: 2367–2379

10. Viotti M, Nowotschin S, Hadjantonakis AK (2014) SOX17 links gut endoderm morphogenesis and germ layer segregation. Nat Cell Biol 16: 1146–1156

11. Ibarra-Soria X et al (2018) Defining murine organogenesis at single-cell resolution reveals a role for the leukotriene pathway in regulating blood progenitor formation. Nat Cell Biol 20: 127–134

12. Nowotschin S, Hadjantonakis AK (2018) Lights, Camera, Action! Visualizing the Cellular Choreography of Mouse Gastrulation. Dev Cell 47: 684–685

13. Warmflash A, Sorre B, Etoc F, Siggia ED, Brivanlou AH (2014) A method to recapitulate early embryonic spatial patterning in human embryonic stem cells. Nat Methods 11: 847–854

14. van den Brink SC, Baillie-Johnson P, Balayo T, Hadjantonakis AK, Nowotschin S, Turner DA, Martinez Arias A (2014) Symmetry breaking, germ layer specification and axial organisation in aggregates of mouse embryonic stem cells. Development 141: 4231–4242

15. Turner DA et al (2017) Anteroposterior polarity and elongation in the absence of extra-embryonic tissues and of spatially localised signalling in gastruloids: mammalian embryonic organoids. Development 144: 3894–3906

16. Beccari L, Moris N, Girgin M, Turner DA, Baillie-Johnson P, Cossy AC, Lutolf MP, Duboule D, Arias AM (2018) Multi-axial self-organization properties of mouse embryonic stem cells into gastruloids. Nature 562: 272–276

17. Boxman J, Sagy N, Achanta S, Vadigepalli R, Nachman I (2016) Integrated live imaging and molecular profiling of embryoid bodies reveals a synchronized progression of early differentiation. Sci Rep 6: 31623

18. Harrison SE, Sozen B, Christodoulou N, Kyprianou C, Zernicka-Goetz M (2017) Assembly of embryonic and extraembryonic stem cells to mimic embryogenesis in vitro. Science 356:

19. Sozen B et al (2018) Self-assembly of embryonic and two extra-embryonic stem cell types into gastrulating embryo-like structures. Nat Cell Biol 20: 979–989

20. Rivron NC et al (2018) Blastocyst-like structures generated solely from stem cells. Nature 557: 106–111

21. Morgani SM, Metzger JJ, Nichols J, Siggia ED, Hadjantonakis AK (2018) Micropattern differentiation of mouse pluripotent stem cells recapitulates embryo regionalized cell fate patterning. Elife 7:

22. Kubo A, Shinozaki K, Shannon JM, Kouskoff V, Kennedy M, Woo S, Fehling HJ, Keller G (2004) Development of definitive endoderm from embryonic stem cells in culture. Development 131: 1651–1662

23. Acloque H, Adams MS, Fishwick K, Bronner-Fraser M, Nieto MA (2009) Epithelial-mesenchymal transitions: the importance of changing cell state in development and disease. J Clin Invest 119: 1438–1449

24. Kalluri R, Weinberg RA (2009) The basics of epithelial-mesenchymal transition. J Clin Invest 119: 1420–1428

25. Ferrer-Vaquer A, Viotti M, Hadjantonakis AK (2010) Transitions between epithelial and mesenchymal states and the morphogenesis of the early mouse embryo. Cell Adh Migr 4: 447–457

26. Camus A, Perea-Gomez A, Moreau A, Collignon J (2006) Absence of Nodal signaling promotes precocious neural differentiation in the mouse embryo. Dev Biol 295: 743–755

27. Zhou X, Sasaki H, Lowe L, Hogan BL, Kuehn MR (1993) Nodal is a novel TGF-beta-like gene expressed in the mouse node during gastrulation. Nature 361: 543–547

28. Engert S, Burtscher I, Liao WP, Dulev S, Schotta G, Lickert H (2013) Wnt/beta-catenin signalling regulates Sox17 expression and is essential for organizer and endoderm formation in the mouse. Development 140: 3128–3138

29. Yasunaga M et al (2005) Induction and monitoring of definitive and visceral endoderm differentiation of mouse ES cells. Nat Biotechnol 23: 1542–1550

30. ten Berge D, Koole W, Fuerer C, Fish M, Eroglu E, Nusse R (2008) Wnt signaling mediates self-organization and axis formation in embryoid bodies. Cell Stem Cell 3: 508–518

31. Kwon GS, Viotti M, Hadjantonakis AK (2008) The endoderm of the mouse embryo arises by dynamic widespread intercalation of embryonic and extraembryonic lineages. Dev Cell 15: 509–520

32. Etoc F, Metzger J, Ruzo A, Kirst C, Yoney A, Ozair MZ, Brivanlou AH, Siggia ED (2016) A Balance between Secreted Inhibitors and Edge Sensing Controls Gastruloid Self-Organization. Dev Cell 39: 302–315

33. Maldonado M, Luu RJ, Ramos ME, Nam J (2016) ROCK inhibitor primes human induced pluripotent stem cells to selectively differentiate towards mesendodermal lineage via epithelial-mesenchymal transition-like modulation. Stem Cell Res 17: 222–227

34. Korostylev A, Mahaddalkar PU, Keminer O, Hadian K, Schorpp K, Gribbon P, Lickert H (2017) A high-content small molecule screen identifies novel inducers of definitive endoderm. Mol Metab 6: 640–650

35. Galli C, Piemontese M, Lumetti S, Ravanetti F, Macaluso GM, Passeri G (2012) Actin cytoskeleton controls activation of Wnt/beta-catenin signaling in mesenchymal cells on implant surfaces with different topographies. Acta Biomater 8: 2963–2968

36. Gadue P, Huber TL, Paddison PJ, Keller GM (2006) Wnt and TGF-beta signaling are required for the induction of an in vitro model of primitive streak formation using embryonic stem cells. Proc Natl Acad Sci U S A 103: 16806–16811

